# Personalized Functional Brain Network Topography Predicts Individual Differences in Youth Cognition

**DOI:** 10.1101/2022.10.11.511823

**Authors:** Arielle S. Keller, Adam R. Pines, Valerie J. Sydnor, Zaixu Cui, Maxwell A. Bertolero, Ran Barzilay, Aaron F. Alexander-Bloch, Nora Byington, Andrew Chen, Gregory M. Conan, Christos Davatazikos, Eric Feczko, Timothy J. Hendrickson, Audrey Houghton, Bart Larsen, Hongming Li, Oscar Miranda-Dominguez, David R. Roalf, Anders Perrone, Sheila Shanmugan, Russell T. Shinohara, Yong Fan, Damien A. Fair, Theodore D. Satterthwaite

## Abstract

Individual differences in cognition during childhood are associated with important social, physical, and mental health outcomes in adolescence and adulthood. Given that cortical surface arealization during development reflects the brain’s functional prioritization, quantifying variation in the topography of functional brain networks across the developing cortex may provide insight regarding individual differences in cognition. We test this idea by defining personalized functional networks (PFNs) that account for interindividual heterogeneity in functional brain network topography in 9-10 year olds from the Adolescent Brain Cognitive Development^SM^ Study. Across matched discovery (n=3,525) and replication (n=3,447) samples, the total cortical representation of fronto-parietal PFNs positively correlated with general cognition. Cross-validated ridge regressions trained on PFN topography predicted cognition across domains, with prediction accuracy increasing along the cortex’s sensorimotor-association organizational axis. These results establish that functional network topography heterogeneity is associated with individual differences in cognition before the critical transition into adolescence.

## INTRODUCTION

Individual differences in cognition during childhood are associated with academic performance^1^ and quality of life in youth,^2^ as well as social, physical and mental health outcomes in adulthood.^3–5^ Moreover, cognitive deficits during youth are associated with heightened risk for psychopathology,^6^ risk-taking behaviors,^7^ cardiovascular disease,^8–10^ and all-cause mortality.^11,12^ Understanding how individual differences in cognitive functioning emerge during childhood is a critical prerequisite for efforts that seek to promote healthy neurocognitive development. Prior neuroimaging studies have demonstrated that complex cognitive tasks engage spatially-distributed, large-scale association networks.^13–15^ However, less is known about the relationship between individual differences in cognition and the spatial layout of functional networks on the anatomic cortex – an individual’s *functional topography*. Attempts at investigating this important problem have faced two key challenges. First, methods must account for person-specific variation in functional topography across individuals, which is especially pronounced in association cortex. Second, recent studies have emphasized that reproducible brain-behavior associations may require very large samples.^17^ We sought to overcome these challenges by capitalizing upon recent advances in machine learning to identify individual-specific functional brain networks in large discovery and replication samples. We tested the overarching hypothesis that the functional topography of networks in association cortex would be associated with individual differences in cognitive function in children.

Studies in humans using fMRI have typically studied functional brain networks using a “one-size-fits-all” approach with standardized network atlases.^18,19^ In this approach, a 1:1 correspondence between structural and functional neuroanatomy across individuals is assumed as fMRI data is co-registered to a structural image, and then normalized to a structural template. This critical assumption has been proven to be demonstrably false by studies from multiple independent laboratories.^20–23^ These studies have revealed substantial inter-individual heterogeneity in functional topography,^20–25^ with especially notable heterogeneity in networks in association cortex that support higher-order cognition.^21^ To overcome this challenge, precision functional mapping techniques have been developed as an alternative to using group-level atlases. These techniques are used to derive individually-defined networks that capture each brain’s unique pattern of functional topography. Such personalized functional networks (PFNs) have been found to be highly stable within individuals and to predict an individual’s spatial pattern of activation on fMRI tasks.^21,22,26^

Notably, the same networks that both support higher-order cognition and have the greatest variability in functional topography tend to lie near the upper end of a predominant axis of hierarchical cortical organization known as the sensorimotor-association (S-A) axis, which spans from unimodal visual and somotomotor cortex to transmodal association cortex.^29^ The S-A axis summarizes the canonical spatial patterning of numerous cortical properties, including myelination, evolutionary expansion, transcriptomics, metabolism, and the principal gradient of functional connectivity.^30^ Prominent individual variation in the functional topography of networks at the association pole—including the frontoparietal network, ventral attention network, and default mode network—has been posited to impact individual differences in cognition.^23^ Indeed, our collaborative group^16^ recently reported that greater total cortical representation of fronto-parietal PFNs was associated with better cognitive performance, and found that a model trained on the complex pattern of association network functional topography could predict cognition in unseen data. However, while these results were drawn from a large study, it was collected at a single site, and has not yet been replicated. This limitation points to the ongoing challenge of reproducibility in studies that seek to define brain-behavior relationships in humans. The reproducibility crisis has been documented extensively,^33,34^ marked by failed replications of high-profile findings,^35,36^ and prompted a renewed emphasis on methods to increase the generalizability of computational models to new datasets.^37^ In addition to the well-documented problems arising from small sample sizes^17^ and over-fitting,^38^ it may also be the case that a lack of consideration for individual-specific neuroanatomy has also contributed to weak effect sizes and non-reproducible findings of prior work.

We sought to delineate the relationship between functional topography and individual differences in cognition by conducting a replication and extension of Cui et al.^16^ in two large, matched samples of youth from the Adolescent Brain Cognitive Development^SM^ (ABCD) Study^39–41^ (total *n*=6,972). Using spatially-regularized non-negative matrix factorization,^80^ we identified personalized functional brain networks that captured inter-individual heterogeneity in functional topography while maintaining interpretability. We sought to replicate two key results.^16^ First, we sought to replicate the finding that fronto-parietal PFN topography is associated with individual differences in cognition. Second, we aimed to demonstrate that predictive models trained on PFN topography could predict youth cognition in unseen data. Furthermore, we extended prior work and investigated whether PFN topography was predictive of the ability to perform specific cognitive tasks or more broadly associated with general cognitive abilities, by training models to predict three major domains of cognition^42^ (general cognition, executive function, and learning/memory). Finally, we predicted that the strength of associations between functional topography and cognition would align with the cortical hierarchy defined by S-A axis, with the functional topography of PFNs in association cortex yielding the most accurate predictions of individual differences in cognition. As described below, this study constitutes the largest replication of precision functional mapping in children to date, confirming reproducible brain-behavior associations of individual differences in cognition and demonstrating that these relationships align with a major cortical hierarchy.

## RESULTS

We aimed to understand how individual differences in functional brain network topography relate to individual differences in cognitive functioning in a sample of n=6,972 children aged 9-10 years old from the Adolescent Brain Cognitive Development^SM^ (ABCD) Study. To account for inter-individual heterogeneity in the spatial layout of functional brain networks, we used precision functional mapping to define personalized functional brain networks (PFNs) for each individual. Leveraging a previously-defined group atlas^16^, we used an advanced machine learning method – spatially-regularized non-negative matrix factorization – to identify 17 personalized functional networks within each individual (**Figure 1**). This procedure yielded a set of 17 matrices of network weights across each vertex (soft parcellation; used for analysis) as well as a matrix of non-overlapping networks describing the highest network weight at each vertex (hard parcellation; used primarily for visualization). To determine where each PFN fell along a predominant axis of cortical hierarchical organization, we computed the average sensorimotor-association (S-A) axis rank across the vertices within each PFN using the group-averaged hard parcellation.

**Figure 1.**
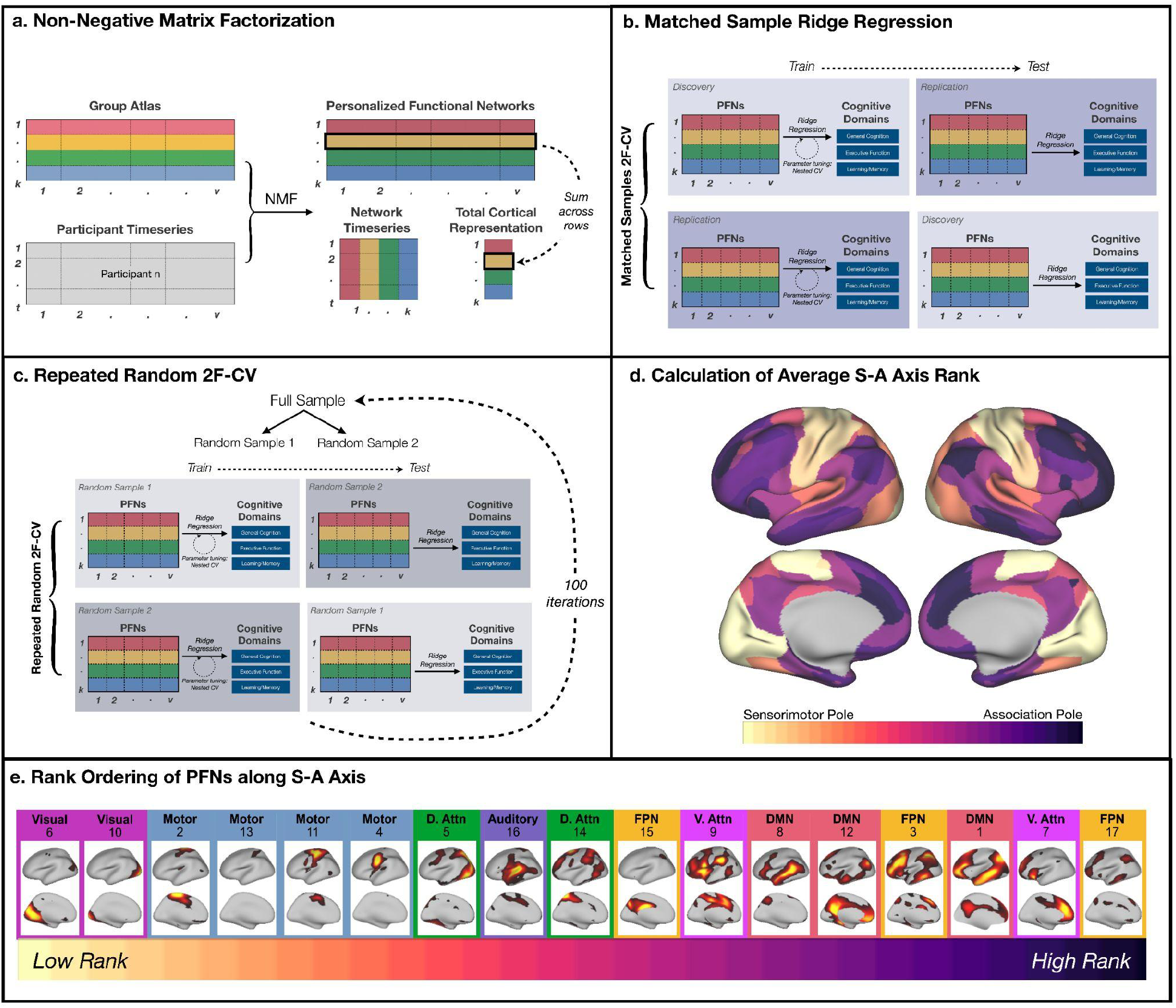
Identification and Analysis of Personalized Functional Brain Networks (PFNs). **a** Using a previously-defined group atlas^16^ as a prior, we generated personalized functional networks (PFNs) by applying NMF to each individual participant’s vertex by time matrix. This procedure allows each network in the consensus group atlas to have a varying cortical representation in each individual, thereby capturing individual differences in the size and layout of networks while simultaneously allowing for interpretable between-individual comparisons. We also calculated the total cortical representation of each PFN by summing each network’s loadings across all vertices. **b** To evaluate whether an individual’s multivariate pattern of PFN topography could accurately predict general cognition in unseen data, we trained linear ridge regression models using the cortical representation of each PFN while controlling for age, sex, site, and head motion. Leveraging our matched discovery and replication samples for out-of-sample testing, we first trained models in the discovery sample using nested cross-validation for parameter tuning, and then tested these models in the held-out replication sample. We then performed nested training in the replication sample and testing in the held-out discovery sample. **c** To confirm that our results were not dependent on the matched discovery and replication sample split, we conducted repeated random cross-validation over one hundred iterations, each time performing a random split of our full sample and applying two-fold cross-validation. **d** Next, we calculated the average sensorimotor-association (S-A) axis rank across the vertices contained within each PFN. **e** We then rank-ordered each PFN according to its average S-A rank. Brain maps depict vertex loadings for each PFN.

### The total cortical representation of fronto-parietal PFNs is associated with cognitive performance

We sought to replicate previously-reported associations between the functional topography of PFNs and cognitive performance.^16^ We previously found that greater cortical representation of two of the three fronto-parietal networks (networks 15 and 17) was associated with better cognition (see Figure 6 in Cui et al.). Here, general cognition was operationalized as the first principal component from a Bayesian probabilistic principal components analysis, capturing the largest amount of variance across nine cognitive tasks;^42^ we hypothesized that general cognition would show stronger associations with functional topography than secondary or tertiary cognitive domains. Notably, while the first cognitive accuracy factor from our prior report is typically referred to as “executive function and complex cognition” (and abbreviated as “executive function”), it most aligns with the general cognition factor from ABCD.^85^

As previously,^16^ we first calculated the total cortical representation of each PFN as the sum of network loadings across all vertices, using the soft parcellation to account for spatial overlap across functional brain networks. We then applied linear mixed-effects models to probe the association between total cortical representation of each PFN and general cognition while accounting for age, sex, family, and head motion (mean FD) as model covariates (**Table 1; Figure 2**); multiple comparisons were accounted for using the Bonferroni method. Note that ComBat harmonization was applied to account for variability across sites.^77,78^ We found that all three fronto-parietal PFNs (networks 3, 15, and 17) were significantly positively associated with general cognition across both the discovery and replication samples. Together, these results replicate the findings presented in Figure 6 of Cui et al.^16^ In addition to replicating these prior results regarding fronto-parietal network topography in both samples, we additionally found that one somatomotor network (network 4) was inversely associated with cognitive performance in both discovery and replication samples, and another somatomotor network (network 2) was inversely associated with cognition in only the discovery sample. Notably, the total cortical representation of network 2 was similarly found to be inversely related to cognition in the original report by Cui et al.^16^

**Table 1.** Linear mixed effects models depicting associations between General Cognition and fronto-parietal PFN topography. Note that data were harmonized across sites using ComBat^77,78^ and each model also included a random effect term for family ID.

| *Predictors* | PFN 3 |  |  |  | PFN 15 |  |  |  | PFN 17 |  |  |  |
| --- | --- | --- | --- | --- | --- | --- | --- | --- | --- | --- | --- | --- |
| | $\beta$ | *Std. Error* | *t* | *p<sub>bonf</sub>* | $\beta$ | *Std. Error* | *t* | *p<sub>bonf</sub>* | $\beta$ | *Std. Error* | *t* | *p<sub>bonf</sub>* |
| **Discovery** |  |  |  |  |  |  |  |  |  |  |  |  |
| Intercept | 0.02 | 0.02 | 0.69 | 0.488 | 0.07 | 0.02 | 2.95 | **0.003** | -0.04 | 0.02 | -1.52 | 0.128 |
| Age | -0.04 | 0.02 | -2.54 | **0.011** | -0.02 | 0.02 | -1.28 | 0.201 | -0.04 | 0.02 | -2.58 | **0.010** |
| Sex | -0.05 | 0.03 | -1.60 | 0.109 | -0.15 | 0.03 | -4.51 | **6.72 x 10<sup>-6</sup>** | 0.05 | 0.03 | 1.57 | 0.117 |
| Mean FD | 0.12 | 0.02 | 6.98 | **3.44 x 10<sup>-12</sup>** | 0.12 | 0.02 | 6.82 | **1.06 x 10<sup>-11</sup>** | 0.04 | 0.02 | 2.22 | **0.027** |
| General Cognition | 0.08 | 0.02 | 3.24 | **0.001** | 0.09 | 0.02 | 3.50 | **4.67 x 10<sup>-4</sup>** | 0.10 | 0.02 | 4.29 | **1.88 x 10<sup>-5</sup>** |
| **Replication** |  |  |  |  |  |  |  |  |  |  |  |  |
| Intercept | 0.01 | 0.02 | 0.43 | 0.665 | 0.07 | 0.02 | 2.82 | **0.005** | -0.02 | 0.02 | -1.00 | 0.320 |
| Age | -0.01 | 0.02 | -0.84 | 0.400 | -0.06 | 0.02 | -3.32 | **9.02 x 10<sup>-4</sup>** | -0.05 | 0.02 | -2.65 | **0.008** |
| Sex | -0.04 | 0.03 | -1.28 | 0.199 | -0.15 | 0.03 | -4.27 | **2.01 x 10<sup>-5</sup>** | 0.04 | 0.03 | 1.11 | 0.265 |
| Mean FD | 0.16 | 0.02 | 9.05 | **2.39 x 10<sup>-19</sup>** | 0.08 | 0.02 | 4.54 | **5.81 x 10<sup>-6</sup>** | 0.04 | 0.02 | 2.11 | **0.035** |
| General Cognition | 0.08 | 0.02 | 3.07 | **0.002** | 0.09 | 0.02 | 3.74 | **1.84 x 10<sup>-4</sup>** | 0.12 | 0.02 | 4.68 | **2.98 x 10<sup>-6</sup>** |

**Figure 2.**
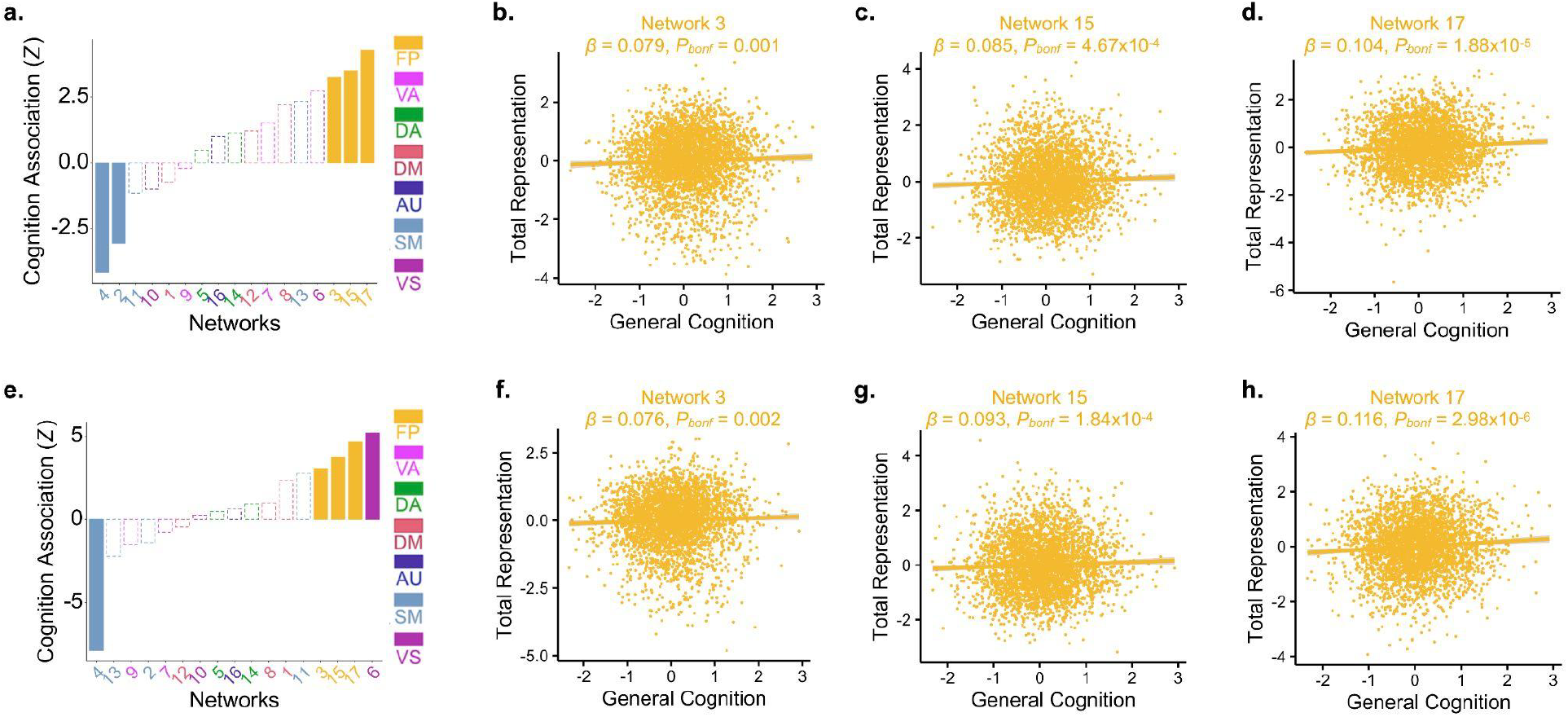
Total cortical representation of fronto-parietal PFNs are positively associated with cognition. Ordering the seventeen PFNs by the strength of their signed association with general cognition, we found significant positive associations between general cognition and the total cortical representation of all three fronto-parietal PFNs and negative correlations with a somatomotor network in both the discovery (**a-d**) and replication (**e-h**) samples (P_Bonf_ < 0.05; dashed lines indicate networks with non-significant effects). Scatterplots depict the relationship between general cognition and the total cortical representation of fronto-parietal networks 3, 15, and 17.

### PFN topography predicts individual differences in cognition

Building on our findings that univariate fronto-parietal PFN topography is positively associated with cognition, we next sought to replicate the prior finding that the multivariate pattern of PFN topography could predict cognitive performance in unseen data. As previously,^16^ we trained ridge regression models using the cortical representation of each PFN (network loadings at each vertex) while controlling for age, sex, site, and head motion. Leveraging our matched discovery and replication samples for out-of-sample testing, we first trained models in the discovery sample using nested cross-validation for parameter tuning, and then used the held-out replication sample for testing. We then performed the opposite procedure, performing nested training in the replication sample and testing in the held-out discovery sample. We found that individualized functional topography accurately predicted out-of-sample cognitive performance in both samples (**Figure 3a**, discovery: *r* = 0.41, *p*<0.001, 95% CI: [0.39, 0.44]; replication: *r* = 0.45, *p*<0.001, 95% CI: [0.43, 0.48]). Confirming that our results were not dependent on the matched discovery and replication sample split, we also applied repeated random cross-validation over one hundred repetitions as previously,^16^ which returned similar results (**Figure 3b**, mean *r* = 0.44, *p*<0.001). These results show remarkably high consistency with correlations between actual and predicted cognitive performance reported in prior work^16^ (Matched sample 1: *r* = 0.46, *p*<0.001; Matched sample 2: *r* = 0.41, *p*<0.001; Repeated random CV: mean *r* = 0.42, *p*<0.001; see Figure 7 in Cui et al.).

**Figure 3.**
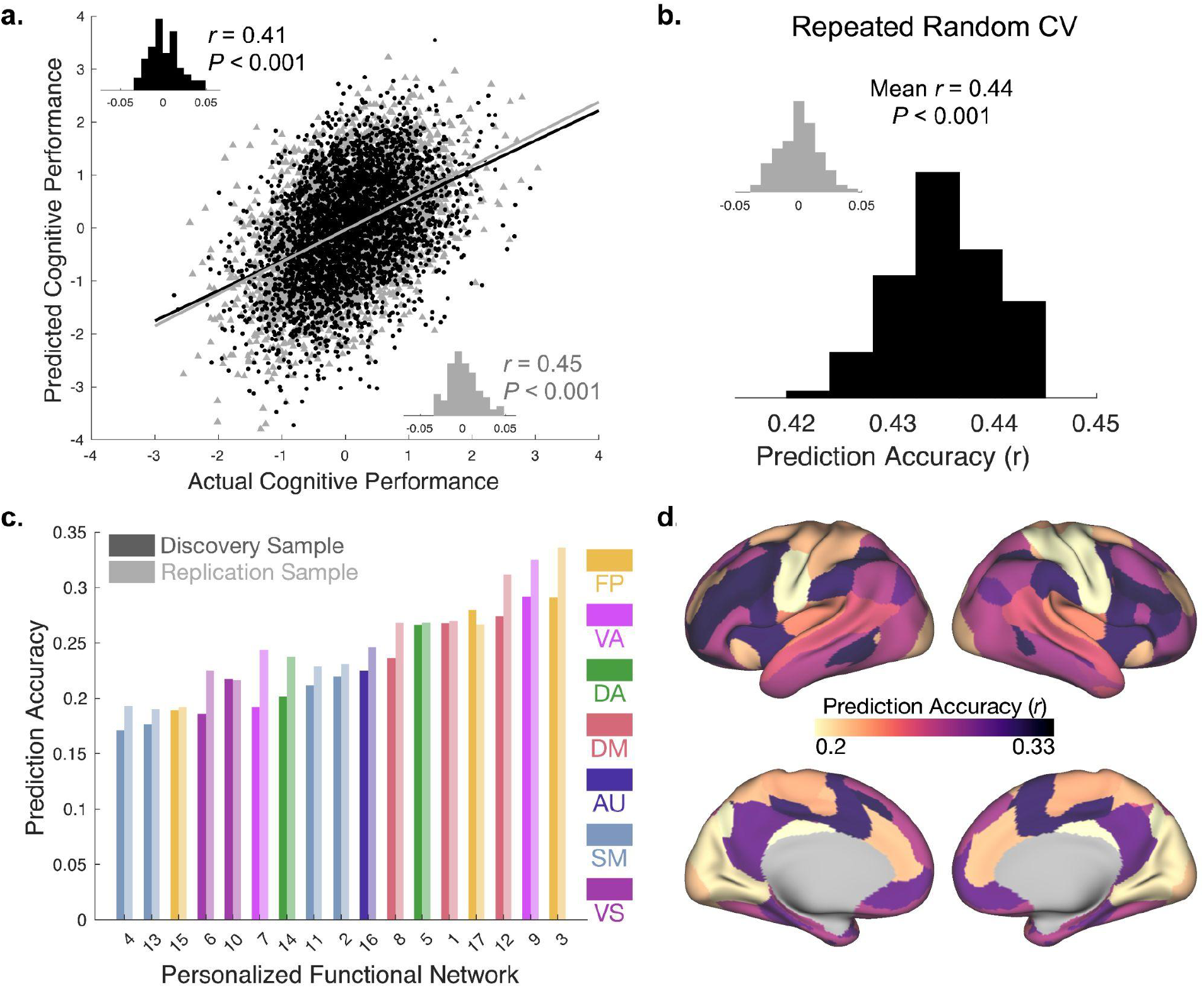
Functional Topography of Association Networks Predicts Individual Differences in General Cognition. **a** Association between actual and predicted cognitive performance using two-fold cross-validation (2F-CV) with nested cross-validation for parameter tuning across both the discovery (black scatterplot) and replication (gray scatterplot) samples. Inset histograms represent the distributions of prediction accuracies from a permutation test. **b** Repeated random 2F-CV (100 runs) provided evidence of stable prediction accuracy across splits of the data, which was far better than a null distribution with permuted data (inset). **c** The topographic features in the 17-PFN model that contributed most to prediction accuracy were found in association cortex critical for higher-order cognition, and were maximal in the ventral attention and fronto-parietal control networks. Note that all *p*-values associated with prediction accuracies are significant after Bonferonni correction for multiple comparisons. **d** Functional topography within the association cortex drives prediction of general cognition. Prediction accuracy across the full sample shown for seventeen cross-validated models trained on each PFN independently.

To evaluate the relative contributions of each network to prediction accuracy, we trained linear ridge regression models on the functional topography of each PFN independently. We found that the fronto-parietal and ventral attention networks tended to have the highest prediction accuracies, whereas the somato-motor and visual networks tended to have the lowest (**Figure 3c**,**d**). These results are again highly consistent with the feature weights from models which used all features and align with our prior report^16^ (see **Supplementary Figure 2** for exact replication), with striking consistency in prediction accuracies across datasets and samples. Together, these results suggest that individual variation in functional network topography has important implications for cognitive performance in childhood.

### PFN topography predicts executive function and learning/memory with reduced accuracy

We next evaluated whether multivariate patterns of PFN topography could be used to predict cognitive performance in held-out data across other cognitive domains. We again trained linear ridge regression models using PFN topography and identical covariates to predict either executive function or learning/memory, which are the second and third ranked principal components capturing variance across nine cognitive tasks.^42^ Although, it is worth noting that while these prediction accuracies were less strong than for the first principal component of general cognition, we found that individualized functional topography predicted performance in our two samples for both executive function (**Figure 4a**, discovery: *r* = 0.17, *p*<0.001, 95% CI: [0.14, 0.20]; replication: *r* = 0.16, *p*<0.001, 95% CI: [0.13, 0.20]) and learning/memory (**Figure 4e**, discovery: *r* = 0.27, *p*<0.001, 95% CI: [0.24, 0.30]; replication: *r* = 0.27, *p*<0.001, 95% CI: [0.24, 0.30]). Repeated random two-fold cross-validation again returned similar results (**Figure 4b**, mean *r* = 0.17, *p*<0.001; **Figure 4f**, mean *r* = 0.28, *p*<0.001). When ridge regression models were trained using the topography of each PFN independently, fronto-parietal and ventral attention PFNs yielded the highest prediction accuracies for both executive function (**Figure 4c**,**d**) and learning/memory (**Figure 4d**,**h**).

**Figure 4.**
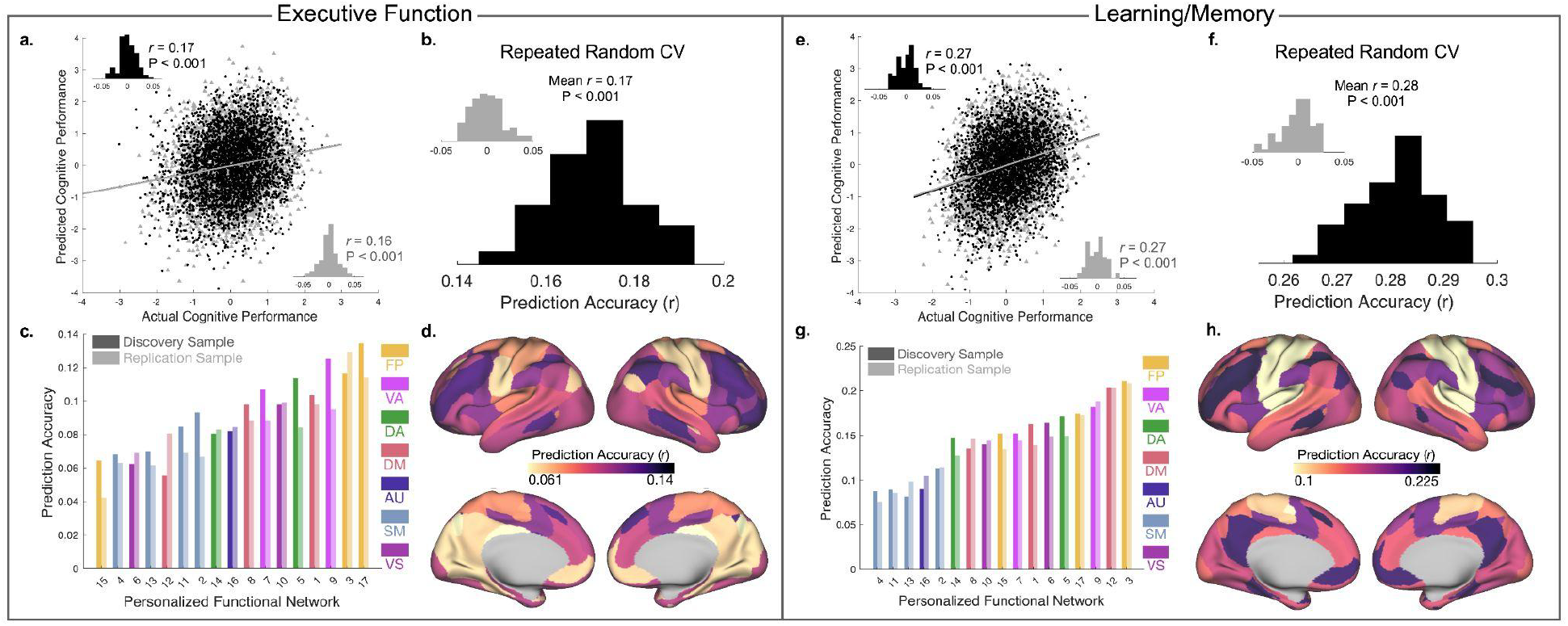
Functional Topography of Association Networks Predicts Individual Differences in Multiple Cognitive Domains. Results of ridge regression models predicting individual differences in executive function (**a-d**) and learning/memory (**e-h**). Panels a/e: Association between actual and predicted executive function (**a**) or learning/memory (**e**) using 2F-CV across both the discovery (black scatterplot) and replication (gray scatterplot) samples. Inset histograms represent the distributions of prediction accuracies from a permutation test. Repeated random 2F-CV (100 runs) provided evidence of stable prediction accuracy across many splits of the data for both executive function (**b**) and learning/memory (**f**), which was far better than a null distribution with permuted data (inset). The PFNs with the highest prediction accuracies for executive function (**c**,**d**) and learning/memory (**g**,**h**) were found in association cortex critical for higher-order cognition, and were maximal in the ventral attention and fronto-parietal control networks. Prediction accuracy shown for seventeen models trained on each PFN independently. Note that all *p*-values associated with prediction accuracies are significant after Bonferonni correction for multiple comparisons.

### Links between functional topography and cognition align with a network’s position in the cortical hierarchy

Motivated by our observation that fronto-parietal association network topography contributed most to the prediction of cognitive performance while somato-motor networks contributed the least, we next investigated whether the predictive accuracy of a given network’s ridge regression model was associated with that network’s rank along the sensorimotor-association (S-A) axis. To account for the spatial auto-correlation of the data, testing used a widely-used spin-based spatial permutation procedure.^43^ We found that prediction accuracy and position along the S-A axis were significantly correlated for predictions of general cognition (Spearman *r*(17)=0.601, *p*_*spin*_=0.012) executive function (Spearman *r*(17)= 0.547, *p*_*spin*_=0.025) and learning/memory (Spearman *r*(17)= 0.537, *p*_*spin*_=0.028; **Figure 5**). These results demonstrate that a network’s position along the S-A axis is associated with the relevance of its functional topography in predicting cognitive performance during childhood).

**Figure 5.**
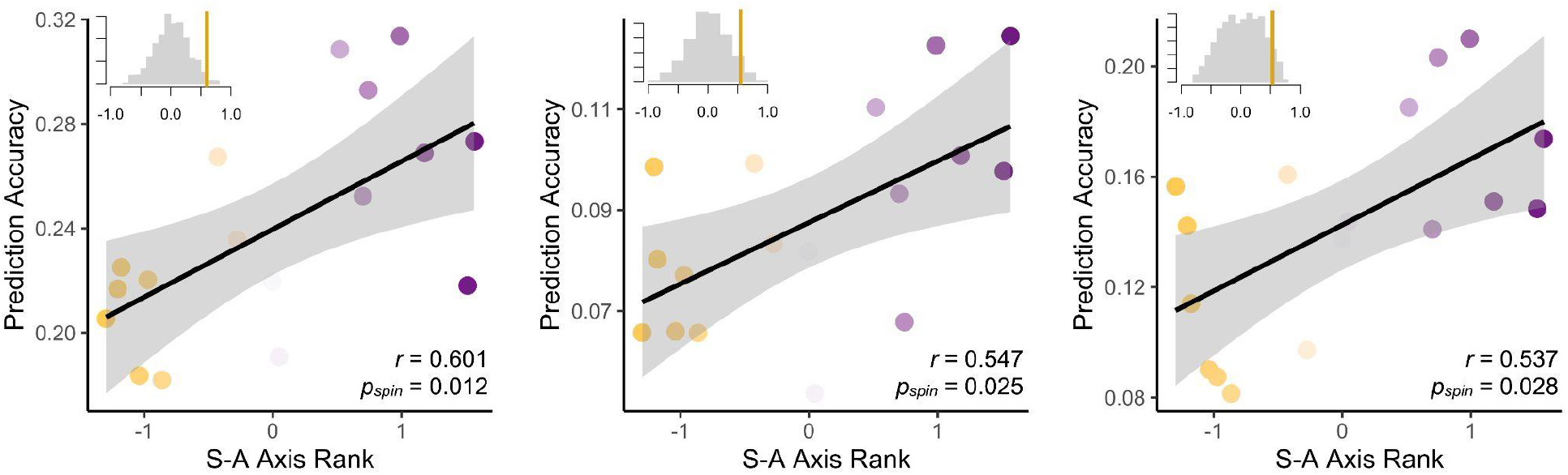
Predictive accuracy of functional topography varies systematically along the S-A axis. The sensorimotor-association (S-A) axis represents a hierarchy of cortical organization. The prediction accuracies of models trained on each PFN independently are significantly associated with the rank of each PFN along the S-A axis across all three cognitive domains: general cognition (left), executive function (middle), and learning/memory (right). Note: average S-A axis ranks for each PFN are z-scored for visualization purposes. Inset histograms depict the distribution of Spearman correlations between rank and prediction accuracy for 1,000 spin-based permutations of the S-A axis, with the vertical line showing the true Spearman correlation value.

## DISCUSSION

In the largest study to use precision functional brain mapping in children to date, we found reproducible associations between individual differences in functional brain network organization and individual differences in cognition. Replicating key findings from a prior study^16^ in samples that were an order of magnitude larger, we confirmed that greater representation of individually-defined fronto-parietal networks is associated with better general cognitive functioning. Furthermore, using cross-validated models trained on the complex multivariate pattern of personalized functional network topography, we were able to predict individual differences in cognitive functioning in unseen participants’ data. Critically, we identified a consistent spatial pattern that accounts for these results, whereby association network topography yields the strongest predictions and sensorimotor network topography yields the weakest predictions of cognitive functioning, directly aligning with the sensorimotor-association (S-A) axis of hierarchical cortical organization.^29^ Together, these findings demonstrate that the link between functional network topography and cognition in children on the precipice of the transition to adolescence is highly reproducible, representing a critical step toward understanding healthy neurocognitive development.

### Scalable Precision Functional Brain Mapping in Children

Our approach successfully overcame two key challenges: addressing inter-individual heterogeneity in functional brain network organization using precision functional mapping and addressing the need for reproducibility by developing cross-validated models in two large samples of thousands of individuals. The reproducibility crisis continues to pose a significant challenge for neuroscience and psychology research, with recent findings further emphasizing the need for very large sample sizes to uncover reproducible brain-wide associations with behavior.^17^ However, large-scale open-source datasets such as the ABCD Study® provide significant hope for a feasible path forward. Our results represent the largest successful replication of associations between functional brain network topography and cognition in children, with remarkably consistent findings across datasets and samples. Several important factors are likely to have contributed to this success. First, both the original study^16^ and our replication leveraged datasets with a previously unprecedented number of fMRI scans of children’s brains (n=693 in the original study, and n=6,972 in our replication). These datasets provided sufficient power to uncover reliable associations between functional brain network topography and cognition. Second, our studies made use of predictive models that were trained and tested using rigorous cross-validation across independent samples. Third, our precision functional brain mapping approach of identifying unique functional networks in individual children’s brains allowed us to capitalize on inter-individual variability rather than treat such variability as noise. This approach may have contributed to the relatively larger effect sizes we observed compared with prior studies using group atlases, which may have bolstered our ability to reproduce these findings. Our scalable precision functional mapping approach may be leveraged in other studies of children and adolescents to harness individual variability and identify important reproducible brain-behavior associations.

### Association network topography supports domain-general cognitive abilities

Having replicated key brain network-cognition associations from prior work,^16^ we can begin to interpret these associations in the context of brain development. Our observation that general cognitive abilities are more strongly associated with PFN topography than other cognitive domains suggests that greater spatial representation of association networks across the cortex may support domain-general cognitive abilities. This finding builds upon prior results highlighting the predominance of general cognitive abilities (also referred to as a “g-factor”^44,45^) in accounting for shared variance across cognitive tasks. Recent work in the ABCD dataset has highlighted the potential role of this g-factor in mediating between genetic risk and psychopathology in children,^46^ suggesting that our identification of functional topography patterns associated with general cognition may represent a brain feature of interest for future studies of resiliency.

We also found a remarkably similar pattern across all three cognitive domains in terms of which PFNs most strongly contributed to predictions of cognitive performance. This consistent pattern was well-described by a major hierarchy of cortical organization known as the S-A axis, with association networks contributing the most to associations with and predictions of cognitive performance across domains. This finding provides further evidence for the existence of a perception-cognition processing hierarchy in the brain^30,47^ that aligns with the S-A axis.^29^ Indeed, the association networks whose topography was most strongly associated with cognitive performance in children also show the greatest evolutionary expansion between non-human primates and humans^48,49^ and their function has been correlated with cognitive performance in adults.^13–15^ Prior studies have also demonstrated that this S-A axis gradually becomes the predominant pattern of cortical functional organizational with age, as the principal gradient of functional connectivity shifts from a visuo-motor axis to the S-A axis from childhood to adolescence^50^—a shift which happens during the protracted development of higher-order cognitive functions.^47^ Future studies may investigate how longitudinal developmental changes in PFN functional topography along the S-A axis is related to the maturation of complex cognitive abilities, complementing cross-sectional work.^16,51^

Another potential explanation for why variability in functional topography in the association cortices is the most strongly associated with individual differences in cognition is that these regions, and particularly regions of the fronto-parietal network, also tend to have the highest degree of inter-individual heterogeneity in other features.^16,19,52–54^ Thus, while various networks across the S-A axis likely *contribute* in diverse ways to cognitive functioning, the notable individual variability in association network topography may be a more salient feature for *predicting* individual differences in cognition. Indeed, these regions tend to exhibit lower structural and functional heritability^55,56^ and undergo the greatest surface area expansion during development.^48^ Moreover, the extended window of plasticity for these regions compared with other parts of the cortex^57^ renders them more likely to be shaped by an individual’s environments and experiences,^56^ potentially further contributing to their unique spatial patterning across individuals. Encouragingly, this extended window in which association networks remain plastic may also indicate that interventions targeting these systems could be effective in supporting the development of healthy cognition.

### Limitations and Future Directions

This study had several limitations worth noting. First, this study was conducted at a single timepoint, using the baseline cohort from the ABCD Study®. As such, we were able to train models that could predict cognitive performance from functional brain network topography in held-out participants data, but were not able to build predictive models of future changes in cognition within an individual. Our work therefore sets the foundation for future longitudinal studies using the ABCD Study® dataset to identify changes in functional brain network organization during development using the personalized functional brain networks we have identified at this baseline assessment. Second, head motion continues to pose an ongoing challenge for neuroimaging studies,^58–60^ and particularly for studies of children.^58^ We have attempted to mitigate these effects by following best practices for reducing the influence of head motion on our results, including using a top-performing preprocessing pipeline and inclusion of motion as a covariate in all analyses. Third, we used data combined across four fMRI runs, including two where a behavioral task performed during scanning was regressed from the data, in line with prior studies of PFNs,^16,51^ aiming to maximize the amount of high-quality data for our study. Prior studies have shown that variation in functional networks is primarily driven by inter-individual heterogeneity rather than task-related factors and that intrinsic functional networks are similar during task and rest.^53^ Finally, our analyses focused on characterizing the cortical surface topography of functional brain networks and thus did not include analyses of subcortical regions. Future studies may use precision functional brain mapping approaches in subcortical areas^61,62^ to further our understanding of the role of the subcortex in cognitive development.

Critically, it is known that cognitive impairments in adulthood are common across diverse psychiatric illnesses including mood^63,64^ and anxiety^65–67^ disorders and our current first-line pharmacological treatments fail to target these cognitive symptoms.^68,69^ Longitudinal studies of neurocognitive developmental trajectories may therefore also provide a critical link between functional brain organization in childhood and psychiatric illness in adulthood, with the potential to identify individuals at risk for cognitive impairments prior to the onset of psychiatric illness and in advance of treatment attempts that are likely to fail. Moreover, this study investigated PFNs in 9-10 year old children prior to the transition to adolescence; these children will be followed longitudinally into adulthood as part of the ABCD Study®. These results therefore lay a strong foundation for future work to uncover how PFNs derived at baseline may predict trajectories of change in cognitive functioning during development as longitudinal data is collected. Such studies may reveal distinct or overlapping neurobiological features that are predictive of future cognitive abilities and whose development tracks with the protracted development of higher-order cognition through childhood and adolescence.

## Conclusion

Together, the findings of this study represent a critical advance in our understanding of the link between individual differences in functional brain network organization and individual differences in cognitive functioning in youth. Further, these results successfully replicated prior findings^16^ across two large samples of youth, providing compelling evidence that these observations are generalizable to new samples. Individual differences in cognition in youth are associated with critical physical, mental, social, and educational outcomes in adolescence and adulthood, ranging from academic achievement and financial success to psychopathology, risk-taking behaviors, and cardiovascular disease.^8–10^ Thus, our findings may inform studies that seek to develop interventions that could promote healthy neurocognitive development. By identifying personalized functional brain networks whose functional topography is associated with cognition, we provide a foundation for future longitudinal studies of neurocognitive development and psychopathology.

## MATERIALS AND METHODS

### Participants

Data were drawn from the Adolescent Brain Cognitive Development^SM^ (ABCD) study^39^ baseline sample from the ABCD BIDS Community Collection (ABCC, ABCD-3165^40^), which included *n*=11,878 children aged 9-10 years old and their parents/guardians collected across 21 sites. Inclusion criteria for this study included being within the desired age range (9-10 years old), English language proficiency in the children, and having the ability to provide informed consent (parent) and assent (child). Exclusion criteria included the presence of severe sensory, intellectual, medical or neurological issues that would have impacted the child’s ability to comply with the study protocol, as well as MRI scanner contraindications. As depicted in **Supplementary Figure 1**, we additionally excluded participants with incomplete data or excessive head motion, yielding a final sample of *n*= 6,972.

To test the generalizability of our results, we repeated each of our analyses in both a discovery sample (*n*=3,525) and a separate replication sample (*n*=3,447) that were matched across multiple socio-demographic variables including age, sex, site, ethnicity, parent education, combined family income, and others.^40,41^ Socio-demographic characteristics of participants in the discovery and replication samples may be found in **Table 2**. Nonsignificant differences between participants in the discovery and replication samples were present across any socio-demographic variables, nor were there any significant differences in scores across the three cognitive domains.

**Table 2.**
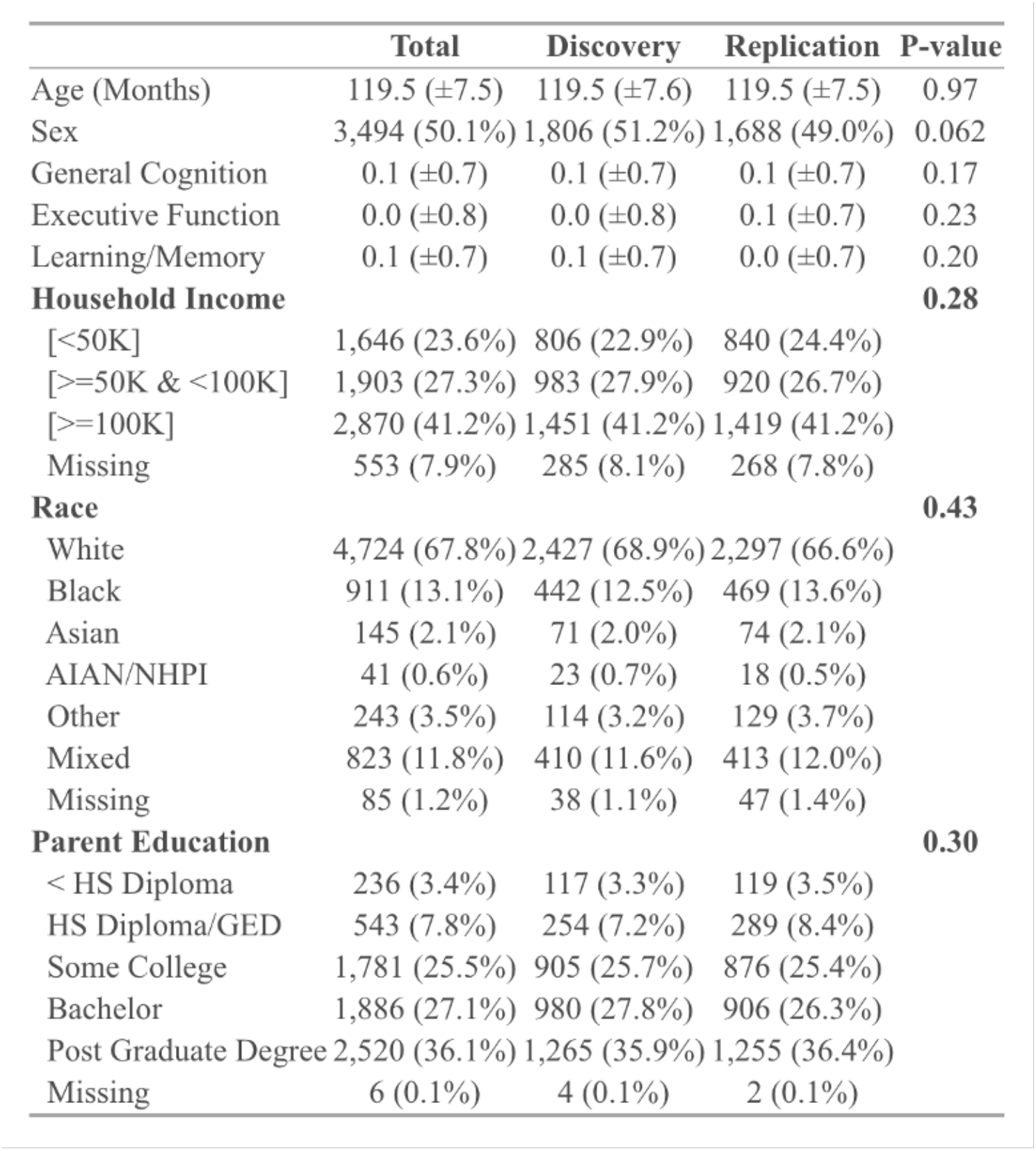
Demographic characteristics and variables of interest in the matched discovery (*n*=3,525) and replication (*n*=3,447) samples. Acronyms: AIAN = American Indian/Alaska Native; NHPI = Native Hawaiian and other Pacific Islander; HS = High School; GED = General Educational Development.

### Cognitive Assessment

Participants completed a battery of cognitive assessments, including seven tasks from the NIH Toolbox (Picture Vocabulary, Flanker Test, List Sort Working Memory Task, Dimensional Change Card Sort Task, Pattern Comparison Processing Speed Task, Picture Sequence Memory Task, and the Oral Reading Test)^70^ as well as two additional tasks (the Little Man Task and the Rey Auditory Verbal Learning Task).^71^ To reduce the dimensionality of these measures and focus our analyses on cognitive domains that explained the majority of behavioral variance in these tasks, we used scores in three previously-established cognitive domains derived from a prior study in this same dataset^42^: 1) general cognition, 2) executive function, and 3) learning/memory. In this study, a three-factor Bayesian Probabilistic Principal Components Analysis (BPPCA) model was applied to the aforementioned battery of nine cognitive tasks. Scores generated by varimax rotated loadings for this three-factor model for general cognition (highest loadings: Oral Reading Test, Picture Vocabulary, and Little Man Task), executive function (highest loadings: Pattern Comparison Processing Speed Task, Flanker Test, and Dimensional Change Card Sort Task), and learning/memory (highest loadings: Picture Sequence Memory Task, Rey Auditory and Verbal Learning Task, and List Sort Working Memory Task) were downloaded directly from the ABCD Data Exploration and Analysis Portal (DEAP).

### Image Processing

Imaging acquisition for the ABCD Study® has been described elsewhere.^72^ As previously described,^40^ the ABCC Collection 3165 from which we drew our data was processed according to the ABCD-BIDS pipeline. This pipeline includes distortion correction and alignment, denoising with Advanced Normalization Tools (ANTS^73^), FreeSurfer^74^ segmentation, surface registration, and volume registration using FSL FLIRT rigid-body transformation.^75,76^ Processing was done according to the DCAN BOLD Processing (DBP) pipeline which included the following steps: 1) de-meaning and de-trending of all fMRI data with respect to time; 2) denoising using a general linear model with regressors for signal and movement; 3) bandpass filtering between 0.008 and 0.09 Hz using a 2nd order Butterworth filter; 4) applying the DBP respiratory motion filter (18.582 to 25.726 breaths per minute), and 5) applying DBP motion censoring (frames exceeding an FD threshold of 0.2mm or failing to pass outlier detection at +/-3 standard deviations were discarded).

Following preprocessing, we concatenated the time series data for both resting-state scans and three task-based scans (Monetary Incentive Delay Task, Stop-Signal Task, and Emotional N-Back Task) as in prior work^16^ to maximize the available data for our analyses. Participants with fewer than 600 remaining TRs after motion censoring or who failed to pass ABCD quality control for their T1 or resting-state fMRI scan were excluded. We additionally excluded participants with incomplete data for our analyses (Supplementary Figure 1). We then applied ComBat harmonization^77,78^ using the *neuroCombat* package protecting age, family and sex as covariates, separately in the discovery and replication samples to harmonize the data across collection sites. Note that for our ridge regression models (described below), we chose to include data collection site as a covariate rather than apply Combat harmonization to avoid leakage across our samples.

### Regularized Non-Negative Matrix Factorization

As previously described,^16,79^ we used non-negative matrix factorization (NMF)^80^ to derive individualized functional networks. NMF identifies networks by positively weighting connectivity patterns that covary, leading to a highly specific and reproducible parts-based representation.^80,81^ Our approach was enhanced by a group consensus regularization term derived from previous work in an independent dataset^16^ that preserves the inter-individual correspondence, as well as a data locality regularization term that makes the decomposition robust to imaging noise, improves spatial smoothness, and enhances functional coherence of the subject-specific functional networks (see Li et al.^79^ for details of the method; see also: https://github.com/hmlicas/Collaborative_Brain_Decomposition). As NMF requires nonnegative input, we re-scaled the data by shifting time courses of each vertex linearly to ensure all values were positive.^79^ As in prior work, to avoid features in greater numeric ranges dominating those in smaller numeric range, we further normalized the time course by its maximum value so that all the time points have values in the range of [0, 1]. For this study, we used identical parameters settings as in prior validation studies,^79^ with the exception of an increase in the data locality regularization term from 10 to 300 to account for smaller vertices in fslr compared with fsaverage5.

### Defining individualized networks

To facilitate group-level interpretations of individually-defined PFNs, we used a group consensus atlas from a previously published study in an independent dataset^16^ as an initialization for individualized network definition. In this way, we also ensured spatial correspondence across all subjects. This strategy has also been applied in other methods for individualized network definition.^23,82^ Details regarding the derivation of this group consensus atlas can be found in previous work.^16^ Briefly, group-level decomposition was performed multiple times on a subset of randomly selected subjects and the resulting decomposition results were fused to obtain one robust initialization that is highly reproducible. Next, inter-network similarity was calculated and normalized-cuts^83^ based spectral clustering method was applied to group the PFNs into 17 clusters. For each cluster, the PFN with the highest overall similarity with all other PFNs within the same cluster was selected as the most representative. The resulting group-level network loading matrix *V* was transformed from fsaverage5 space to fslr space using Connectome Workbench,^84^ and thus the resultant matrix had 17 rows and 59,412 columns. Each row of this matrix represents a functional network, while each column represents the loadings of a given cortical vertex.

Using the previously-derived group consensus atlas^16^ as a prior to ensure inter-individual correspondence, we derived each individual’s specific network atlas using NMF based on the acquired group networks (17 × 59,412 loading matrix) as initialization and each individual’s specific fMRI times series. See Li et al.^79^ for optimization details. This procedure yielded a loading matrix *V* (17 × 59,412 matrix) for each participant, where each row is a PFN, each column is a vertex, and the value quantifies the extent each vertex belongs to each network. This probabilistic (soft) definition can be converted into discrete (hard) network definitions for display and comparison with other methods^19,23,82^ by labeling each vertex according to its highest loading.

### Calculation of Sensorimotor-Association Axis Rank

To compute S-A axis rank for each PFN independently, we computed the average S-A rank across vertices for each PFN according to the hard network parcellation. Original S-A axis ranks by vertex represent the average cortical hierarchy across multiple brain maps^16^ and were derived from https://github.com/PennLINC/S-A_ArchetypalAxis.

### Statistical Analyses

#### Linear Mixed-Effects Models

We used linear mixed effects models (implemented with the “lme4” package in *R*) to assess associations between PFN topography and performance in each cognitive domain while accounting for both fixed and random predictors. All models included fixed effects parameters for age, biological sex, head motion (mean fractional displacement), as well as random intercepts for family (accounting for siblings) and site groupings.

#### Ridge Regression

We trained ridge regression models to predict cognitive performance in each of the three cognitive domains (general cognition, executive function, and learning/memory) using the functional topography (vertex-wise network loading matrices) of each participant’s PFNs. Primary analyses were conducted on models trained on concatenated network loading matrices across the 17 PFNs. Independent network models were also trained on the network-wise loadings at each vertex. All models included covariates for age, sex, site, and motion (mean FD).

Our primary ridge regression models were trained and tested on the ABCD reproducible matched samples^40,41^ using two-fold cross-validation (2F-CV). To ensure that this sample selection procedure did not bias our results, we performed repeated random cross-validation over 100 iterations, each time randomly splitting the sample and repeating the 2F-CV procedure to generate a distribution of prediction accuracies for each model. Furthermore, we used permutation testing to generate null distributions for both the primary models and the repeated random cross-validation models by randomly shuffling the outcome variable. **Supplementary Figure 2** depicts the sum of model weights by PFN for the primary ridge regression models in each of the matched samples.

#### Associations Between Prediction Accuracy and S-A Axis Rank

To compute associations between the prediction accuracy of each individual network model and the average S-A rank for each network, we used Spearman’s rank correlations for each of the three cognitive domains: general cognition, executive function, and learning/memory. To determine whether the alignment between these two spatial maps were driven specifically by the S-A axis, we used spatial permutation testing^43^ (Spin Tests; https://github.com/spin-test/spin-test). The spin test is a spatial permutation method based on angular permutations of spherical projections at the cortical surface. Critically, the spin test preserves the spatial covariance structure of the data, providing a more conservative and realistic null distribution than randomly shuffling locations. With this approach, we applied 1,000 random rotations to spherical representations of S-A axis rank across the cortical surface, each time re-computing the average S-A rank within each PFN and calculating Spearman correlations between prediction accuracy and the permuted average S-A rank in each PFN to generate a null distribution. We then compared the true Spearman correlation value to the null distribution of spatially permuted Spearman correlations by rank ordering.

## Acknowledgment

This study was supported by grants from the National Institute of Health: R01MH113550 (TDS & DSB), R01MH120482 (TDS), R01EB022573 (TDS), R37MH125829 (TDS & DAF), K99MH127293 (BL), F31MH123063-01A1 (AP), R01MH123550 (RTS), R01MH112847 (RTS), R01MH123563 (RTS). ASK was supported by a Neuroengineering and Medicine T32 Fellowship from the National Institute of Neurological Disorders and Stroke (5T32NS091006-08). SS was supported by R25MH119043 and T32NS091008. Additional support was provided by the Penn-CHOP Lifespan Brain Institute.

Data used in the preparation of this article were obtained from the Adolescent Brain Cognitive Development^SM^ (ABCD) Study (https://abcdstudy.org), held in the NIMH Data Archive (NDA). This is a multisite, longitudinal study designed to recruit more than 10,000 children age 9-10 and follow them over 10 years into early adulthood. The ABCD Study® is supported by the National Institutes of Health and additional federal partners under award numbers U01DA041048, U01DA050989, U01DA051016, U01DA041022, U01DA051018, U01DA051037, U01DA050987, U01DA041174, U01DA041106, U01DA041117, U01DA041028, U01DA041134, U01DA050988, U01DA051039, U01DA041156, U01DA041025, U01DA041120, U01DA051038, U01DA041148, U01DA041093, U01DA041089, U24DA041123, U24DA041147. A full list of supporters is available at https://abcdstudy.org/federal-partners.html. A listing of participating sites and a complete listing of the study investigators can be found at https://abcdstudy.org/consortium_members/. ABCD consortium investigators designed and implemented the study and/or provided data but did not necessarily participate in the analysis or writing of this report. This manuscript reflects the views of the authors and may not reflect the opinions or views of the NIH or ABCD consortium investigators. The ABCD data repository grows and changes over time. The ABCD data used in this report came from [NIMH Data Archive Digital Object Identifier 10.15154/1523041]. DOIs can be found at https://nda.nih.gov/abcd.

## Supplemental Figures

**Supplementary Figure 1.**
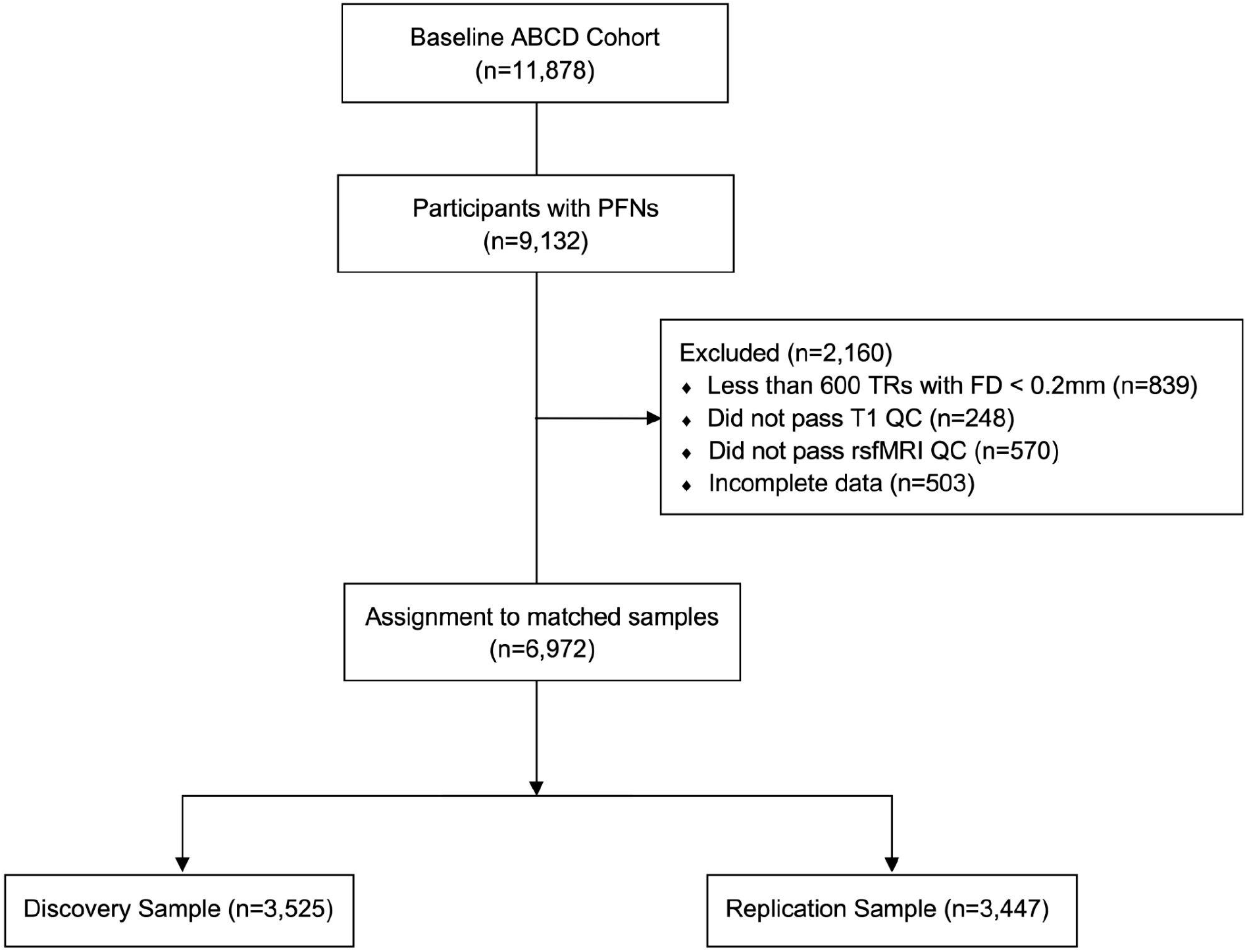
Flow diagram depicting data inclusion and exclusion. Data were drawn from the Adolescent Brain and Cognitive Development (ABCD) study^39^ baseline sample from the ABCD BIDS Community Collection (ABCC, ABCD-3165^40^), which included *n*=11,878 children between the ages of 9-11 years old. Participants were excluded for having incomplete data or excessive head motion, then split into a discovery sample (*n*=3,525) and a matched replication sample (*n*=3,447).^40,41^

**Supplementary Figure 2.**
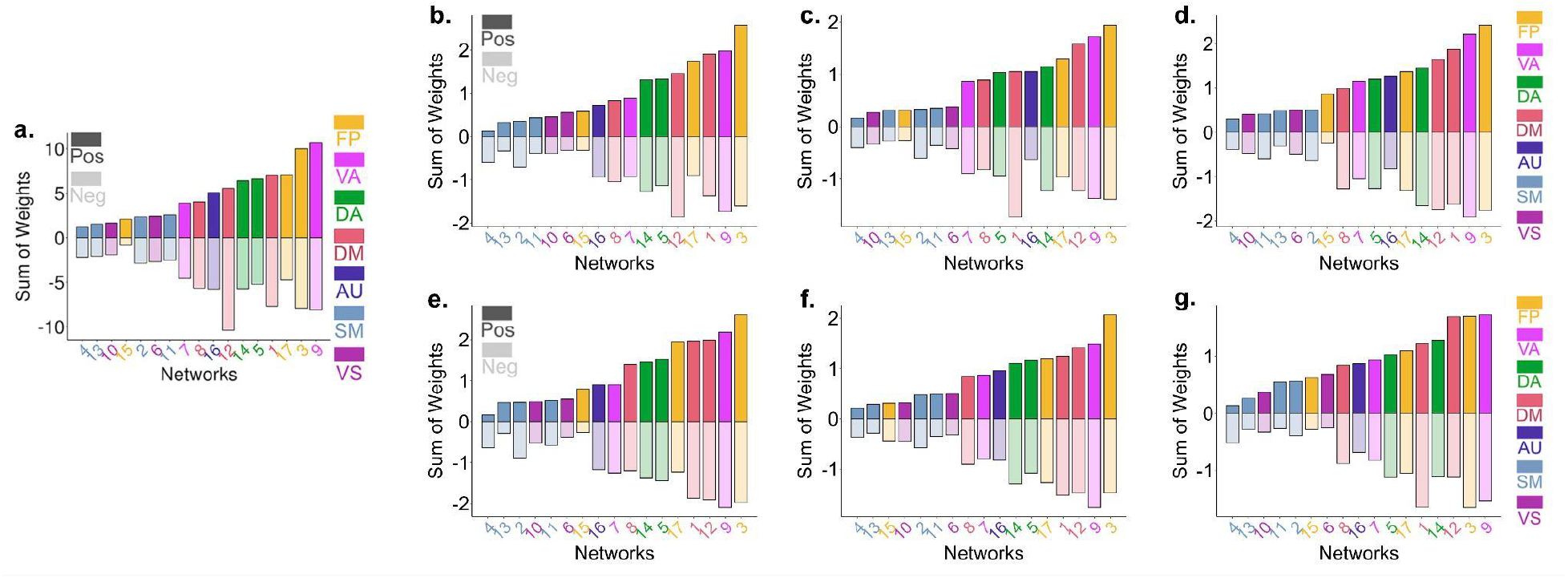
Sum of ridge regression weights by PFN. Replication of results from Cui et al.^31^ (**a**) showing that the PFNs contributing the strongest weights in ridge regression models predicting cognition tend to lie at the associative end of the S-A axis. At each location on the cortex, the absolute contribution weight of each network was summed. **b**,**c**,**d** Models trained on the replication sample and tested in the matched discovery sample; **e**,**f**,**g** Models trained on the discovery sample and tested in the matched replication sample. **b**,**e** general cognition; **c**,**f** executive function; **d**,**g** learning/memory.

## Notes

### Competing Interest Statement

Ran Barzilay reports owning stock in Taliaz Health and serving on the scientific boards of Taliaz Health and Zynerba Pharmaceuticals outside the submitted work.

